# Cobalamin production in phototrophic and mixotrophic cultures of nine duckweed species (Lemnaceae)

**DOI:** 10.1101/2023.10.31.564895

**Authors:** Klamann Linda, Dutta Rhishika, Ghazaryan Lusine, Sela-Adler Michal, Khozin-Goldberg Inna, Gillor Osnat

**Affiliations:** Zuckerberg Institute for Water Research, Blaustein Institutes for Desert Research, Ben-Gurion University of the Negev, Israel; Institute for Bio- and Geoscience – IBG-3, Agrosphere, Forschungszentrum Jülich GmbH, Germany; The French Associates Institute for Agriculture and Biotechnology of Drylands, Blaustein Institutes for Desert Research, Ben-Gurion University of the Negev, Israel

## Abstract

To feed the rapidly increasing world population, food production will have to double while the amount of water and arable land remain unchanged. Therefore, we must transition to environmentally sustainable, but just as nutritious diets by 2050. This will require drastic reduction in animal-based foods and increasing plant-based foods. Plant- based diets are more sustainable and healthier, with but one exception, plants are low in cobalamin (vitamin B_12_). Only a few plant and fungi species contain cobalamin, and some of the cobalamin forms present in algae and mushrooms are not available to humans. Recently, the smallest known plant, the edible duckweed *Wolffia globosa* Mankai, was shown to contain high concentrations of bioavailable cobalamin. We hypothesized that the production of bioavailable cobalamin is not unique to *W. globosa* Mankai but is common to other duckweed species. We also hypothesized that cobalamin production depends on the conditions under which the duckweed is grown. To test our hypotheses, we cultivated nine duckweed species under mixotrophic and phototrophic conditions, then measured the concentration and identified the bioavailability of cobalamin using biological and analytical methods. Our results showed that all duckweed species tested produce bioavailable cobalamin while pseudocobalamin (the non-available form of cobalamin) was not detected. We also found that mixotrophic conditions enhanced duckweed growth but reduced cobalamin concentrations. Our results establish duckweed species as a novel source of plant-derived bioavailable cobalamin and highlight the effect of growth conditions on production. More research is needed to identify the bacteria responsible for the production of cobalamin in duckweed and to optimize the conditions for cobalamin production.

## Introduction

Cobalamin, also known as vitamin B_12_, is an essential vitamin for microorganisms and animals. Cobalamin is involved in normal cell growth and development, acts as a coenzyme that promotes normal carbohydrate and lipid metabolism, and is crucial for DNA synthesis, methylation, and mitochondrial metabolism ^1–3^. Humans require 1-2 µg cobalamin per day, which must be ingested through our diet ^1,2,4^. Long-term B_12_ deficiency can cause serious physiological problems, such as cardiometabolic diseases, hypertension, dyslipidemia, or hyperglycemia ^5^.

Cobalamin is the most widely recognized member of a diverse family of corrinoid cofactors ^6^ that can be found in four different biologically active forms: 5’ deoxyadenosylcobalamin, hydroxocobalamin, methylcobalamin, and cyanocobalamin. The first three compounds naturally occur in food, while cyanocobalamin is the most stable form of cobalamin ^7^. Cobalamin can be formed via aerobic or anaerobic pathways that involve 30 different enzyme-mediated steps ^8,9^. This vitamin can only be produced by bacteria and archaea and was found mainly in tissues of mammals that have ingested foods containing cobalamin or harbor cobalamin-producing bacteria ^6^. Non-animal sources of cobalamin include green macroalgae, mushrooms, fermented foods, synthetic supplements, and ‘natural’ supplements such as the cyanobacteria *Spirulina* and the green microalga *Chlorella* ^10^. However, 83% of the cobalamin in *Spirulina* was found to be pseudocobalamin, a cobalamin analog that is barely absorbed in the mammalian intestine and therefore biologically unavailable to humans ^3,11^. In contrast, the bioavailable form of cobalamin was detected in *Chlorella* products, but fluctuations in pseudocobalamin concentrations were reported ^11^. Pseudocobalamin can block the metabolism of bioavailable cobalamin in mammalian cells ^11^, affecting the bioavailability of cobalamin in *Chlorella*. The unreliable presence of bioavailable cobalamin in algae-based food supplements highlights the importance of detecting a reliable and consistent source of cobalamin in plant-based food sources.

Most plants neither produce nor require cobalamin mostly because they have an alternative methionine synthesis. Yet recently, the duckweed *Wolffia globosa* Mankai was reported to contain 2.8 ± 0.5 µg cobalamin per 100 g dry weight ^2^. Furthermore, duckweed species of the genus *Wolffia* were found to be highly nutritious, containing large amounts of protein, omega-3 fatty acids, dietary fiber, polyphenols, iron, and several other micronutrients ^12–14^. The nutritional benefits of *W. globosa* Mankai for human health were recently demonstrated ^15–19^, including the benefits of the bioavailable cobalamin ^2,20^. However, it is unknown whether the production of bioavailable cobalamin is shared by other duckweed species or unique to *W. globosa* Mankai.

Duckweeds belong to the *Lemnaceae* family and are floating aquatic plants with global distribution. The high nutritional value, high growth rates, high yields, and easy harvest of duckweed could drive their large-scale application as a source of human food or animal feed. Optimizing cultivation strategies to enrich biomass with essential nutrients such as cobalamin could benefit the use of duckweed in large-scale food and feed production. Interestingly, duckweeds are not limited to phototrophic growth driven by CO_2_ assimilation, but they can also utilize different organic carbon sources featuring mixotrophic growth ^21,22^. Different trophic modes have been reported to affect nutrient production, such as starch, in the duckweed species *Spirodela polyrhiza* ^22^.

In this study, we explored the prevalence of cobalamin production in duckweed species as well as the type of cobalamin produced. We also explored the effect of trophic conditions on cobalamin production in the investigated duckweed species. We hypothesized that the production of bioavailable cobalamin, but not pseudocobalamin, is not unique to *W. globosa* Mankai but is a shared trait among members of the duckweed family. We further hypothesized that mixotrophic growth would not only change the growth rate of duckweed ^23^ but also produce different amounts of cobalamin compared to phototrophic growth. To test our hypotheses, we cultivated nine species of duckweed, representing four genera, in controlled environments under photoautotrophic or mixotrophic conditions, at stable pH and nutrient levels. The identification and quantification of bioavailable cobalamin was performed using analytical and microbial assays to test our hypotheses.

## Materials and Methods

### Plant material and growth conditions

Nine different duckweed species from four genera were selected (Table 1) for the growth experiment. Every duckweed species was represented by a single haplotype. The growth medium contained ½ SH with vitamins (PlantMedia, Chiang Mai, Thailand), 53 µM FeNaEDTA (PhytoTech Labs, Lenexa, KS, USA) and 5 mM MES buffer (Carl Roth, Karlsruhe, Germany). The pH was adjusted to 5.8. The medium was sterilized for 20 min at 121°C. For mixotrophic cultivation, 30 mM sucrose (Sigma-Aldrich, St. Louis, MO, USA) from the filtered sterilized stock solution to the sterilized growth medium. The duckweeds were cultivated in flasks covered with an air-permeable stopper (cotton wool) at 25°C in a 16 h light/ 8 h dark cycle at 200 - 250 μmol photons m^-2^ s^-1^ PAR light intensity.

**Table 1.**
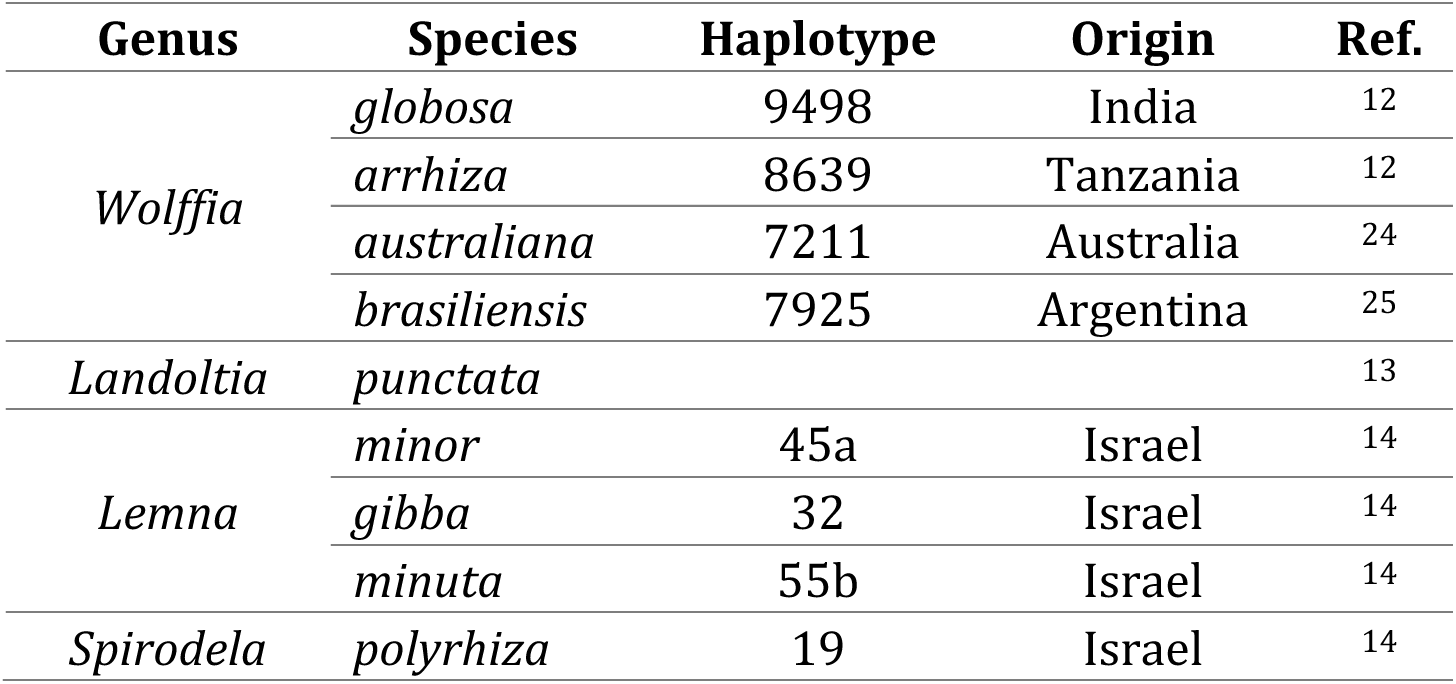
List of the duckweed species used in this study.

| Genus | Species | Haplotype | Origin | Ref. |
| --- | --- | --- | --- | --- |
| *Wolffia* | *globosa* | 9498 | India | 12 |
|  | *arrhiza* | 8639 | Tanzania | 12 |
|  | *australiana* | 7211 | Australia | 24 |
|  | *brasiliensis* | 7925 | Argentina | 25 |
| *Landoltia* | *punctata* |  |  | 13 |
| *Lemna* | *minor* | 45a | Israel | 14 |
|  | *gibba* | 32 | Israel | 14 |
|  | *minuta* | 55b | Israel | 14 |
| *Spirodela* | *polyrhiza* | 19 | Israel | 14 |

Optimal duckweed growth was reported at pH between 5 and 7 ^26^, therefore, the pH of the medium was adjusted throughout the cultivation period to approximately pH 6 by adding 0.1 M NaOH. The growth and photosynthesis of duckweeds were shown to be affected by the availability of organic carbon and nutrients ^23,27^. To test the effect of mixotrophy and nutrient refreshment oncobalaminproduction, four cultivation treatments were applied (Figure 1): (1) mixotrophy + refreshment (cultivation in ½ SH with 30 mM sucrose that was refreshed every 3 - 4 days); (2) mixotrophy (cultivation in ½ SH with 30 mM sucrose); (3) phototrophy + refreshment (cultivation in ½ SH that was refreshed every 3 - 4 days); and (4) phototrophy (cultivation in ½ SH). Rooted duckweed cultures (species of rooted duckweed species of the genera *Lemna, Landoltia,* and *Spirodela*; Table 1) were started with a single frond, while rootless duckweed cultures (species of the genera *Wolffia*; Table 1) were started with five fronds taken from agar plates. For nutrient refreshment, the growth medium was aseptically aspirated and replaced with fresh medium twice a week (every 3 - 4 days).

**Figure 1.**
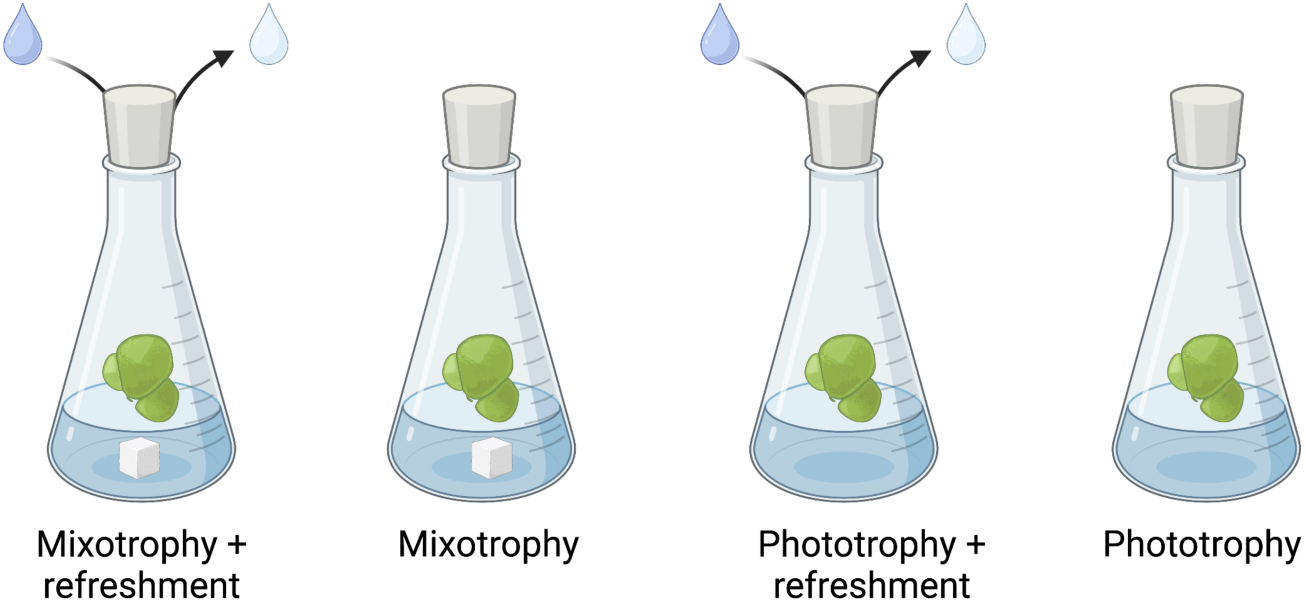
Scheme of growth conditions for the investigation of the cobalamin content in various duckweeds (created with BioRender.com)

Duckweeds were harvested when the surface of the medium in the flask was fully covered. The duckweed fronds were collected using a strainer and dried by shaking for 2 min. The harvested plant was weighed, stored for 24 h at -20°C and then frozen dried by lyophilization (Virtis freeze dryer, ATS Scientific, Warminster, PA, USA) at 4°C.

The relative growth rate (RGR) per day was evaluated as previously described ^28^ with slight modifications (Equation 1, 2)

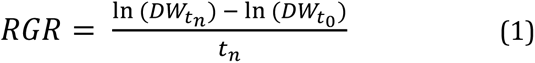

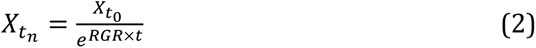

Where RGR have the reciprocal dimension of time as the natural number exponent (Equation 1). Thus, RGR is the increase in parameter value per day value of the duckweed dry weight (DW) at harvest relative to DW at the start of the experiment (t_0_), divided by harvest time (t_n_) in days.

### Chlorophyll analysis

To investigate the growth and photosynthetic performance of cultivated duckweed, chlorophyll a and b pigments were measured. The analysis was carried out as previously described ^29^. Briefly, 1 mL of buffered acetone (80 %) was added to 5 mg of freeze dried duckweed plant material. To avoid chlorophyll transformation into pheophytin due to loss of Mg^2+^, the solution was buffered under neutral or slightly alkaline conditions, therefore, acetone was buffered with HEPEs-KOH buffer (pH 7.4). For homogenization, 1 mm glass beads (Sigma-Aldrich) were added, and the solution was lysed for 1 min using a bead beater (FastPrep homogenizer, MP Biomedicals, Irvine, CA, USA). The lysed solution was incubated in the dark at 4°C overnight and then centrifuged at 10,000 rpm for 5 min to separate the pigments from the plant debris. The supernatant was transferred to a transparent 96-well plate (Corning, Glendale, AZ, USA). The concentration of chlorophyll a and b in the supernatant was measured using a spectrophotometer (Tecan, Männedorf, Switzerland) at 663 and 645 nm, respectively.

Total chlorophyll concentrations per in DW were calculated as previously described ^30,31^ with slight modification using Equations 2-5.

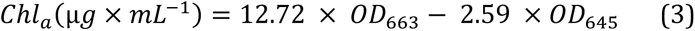

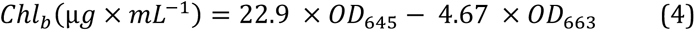

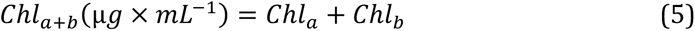

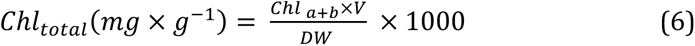

The amount of chlorophyll a and b in the measured volume of duckweed extraction solvent volume (Equations 3-5) was standardized to the duckweed DW by multiplying by the total volume extracted (V) and dividing by the DW of each sample (Equation 6).

### Cobalamin extraction and analysis

For the extraction of cobalamin, 30 – 100 mg of freeze-dried duckweed plant material were weighed and ground to a fine powder using an ULTRA TURRAX (IKA, Staufen, Germany), at 6000 rpm for 2 min. To produce sufficient dry biomass material for the extraction of cobalamin, we had to grow sufficient duckweed biomass at varying growth times for the different species, ranging from three to six weeks. 5 mL of Milli-Q water was added to the grinding tube and the solution was ground again at 6000 rpm for 1 min. Subsequently, the solution was transferred to a 50 mL conical tube, with 3 mL of methanol (100 %). The pH was adjusted by adding 0.5 mL of 0.2 M HCl (pH 4 - 4.5). For the conversion of all forms of cobalamin into the most stable form of cyanocobalamin, 125 µL of KCN (1% w/v; Sigma-Aldrish) was added and the tube was filled to a total of 10 mL with methanol (100 %). The solution was then vortexed and incubated at 90°C for 30 min. Subsequently, the extract was quickly cooled on ice and after cooling to room temperature, the solution was centrifuged at 4000 rpm for 15 min. The supernatant was collected, filtered through a 0.22 µm PVDF filter (MilliporeSigma, Tullagreen, Ireland) and an aliquot of 2 mL was stored at -80°C for MBA analysis and 4.5 mL were further used for solid phase extraction (SPE) C-18 for LC-MS/MS analysis.

### Cobalamin quantification using a microbial bioassay (MBA)

For the determination of total cobalamin in plant samples, a microbiological assay (MBA) was applied. The extracted samples were aliquoted in 2 mL tubes and vaporized under vacuum at 40°C using SpeedVac (Eppendorf, Hamburg, Germany). The samples were resuspended and further diluted in sterile water before applying to MBA. The MBA VitaFast Vitamin B_12_ kit (R-biopharm, Darmstadt, Germany) was conducted according to the manufacturer’s instructions using *Lactobacillus delbrueckii* ssp. *leichmannii* ATCC 7830 as the test microorganism.

### Cobalamin identification and quantification using LC-MS/MS

To detect cobalamin bioavailability (cobalamin and pseudocobalamin) in duckweed extracts, we used a triple quadruple MS/MS Ultivo system with ESI, coupled with an Infinity II 1260 HPLC system (Agilent Technologies, Santa Clara, CA, USA). To purify duckweed extracts, 4.5 mL of the extracted solution was transferred to a glass vial and vaporized with 99.999% N_2_ using TurboVap (Caliper Technology, Hopkinton, MA, USA). The sample was then resuspended in 3 mL of 6.25 % methanol in Milli-Q water. A SPE cartridge C-18 (Phenomenex) was conditioned with 4 mL of methanol (100%) and 4 mL of Milli-Q water, then 3 mL of the sample was loaded. The column was washed with 5 mL acetone (100%) and the analyte was eluted with 5 mL of ethanol and then collected. The elution was vaporized at 50°C with 99.999% N_2_ using TurboVap and resuspended in 300 µL Milli-Q water. Chromatographic separation was applied using a reverse phase C-18 column (Kinetex, Phenomenex, Torrance, CA, USA) by injecting 100 µL of the sample to the column (100 mm ξ 3 mm ξ 2.6 µm) and separation was carried out at 25°C. For mobile phase A, we used 0.1 % aqueous formic acid in Milli-Q water, and for mobile phase B, we used 0.1 % aqueous formic acid in 5 % Milli-Q water and 95 % acetonitrile. The flow rate was 0.5 mL / min using the gradient conditions starting at 10 % of mobile phase B and increasing to 69 % over 5.5 min, then returned to 10 % at 6 min, held for a min and applied post-run of a minute using 10 % mobile phase B. The total run time was 8 min, including the post-run time. Multiple reaction monitoring conditions in a positive ionization mode were used according to Sela et al. (2020).

### Standard solutions

A stock solution of 10 µM cyanocobalamin stock (PhytoTech Labs) stock diluted in methanol (100%) to prepare a standard calibration curve of five points: 0.25, 0.5, 1, 10 and 25 nM. Because pseudocobalamin is not commercially available, we used Spirulina extracts (Spirulina Full Life, Migdal HaÈmek, Israel) to verify the presence of pseudocobalamin, as previously reported ^32^. The solutions were measured by the LC-MS/MS method described above.

### Statistical analysis

All experiments were performed at least three times. Data were tested for the presence of outliers using a Rosener outlier test, and outliers were removed from the data before statistical analysis. For the results of the average growth rate, chlorophyll concentration and cobalamin content were evaluated by two-way ANOVA to compare the samples.

Significant differences between variables were considered if less than 0.01.

## Results

### The effect of nutrient refreshment on the duckweeds

Under both mixotrophic or phototrophic conditions, the supply of fresh nutrients and the adjustment of pH during the cultivation of duckweed cultures (refreshments) did not affect growth, chlorophyl content, or cobalamin accumulation (two- way ANOVA, p > 0.01; Table S1 and S2; Figure S1). Our results suggest that the nutrient content in the ½ SH medium was sufficient throughout the cultivation period (Table S1 and S2; Figure S1). Based on these results, the subsequent analyzes presented in the following ignored the periodic renewal of the cultivation medium.

### Growth rate of duckweed species

The RGR of the duckweed cultures was estimated under mixotrophic and phototrophic conditions. The average RGR when sucrose was added ranged from 0.3 to 0. 5 per day, while the phototrophic RGR ranged from 0.2 to 0.35 per day (Figure 2). Mixotrophic conditions had a significant positive effect on the average growth rate (ANOVA, p < 0.01; Table S1). Furthermore, we have detected a significant difference (ANOVA, p = 0.01; Table S1) between the RGR of rooted species of the genera *Lemna*, *Landoltia*, and *Spirodela* and the rootless species of the genus *Wolffia* cultivated in the presence of sucrose. In contrast, the differences in RGR between the genera cultivated under phototrophic conditions were not significant (ANOVA, p > 0.01; Table S1). Furthermore, both trophic conditions did not result in significant differences in RGR within the rooted species or the rootless species (ANOVA, p > 0.01; Table S1).

**Figure 2.**
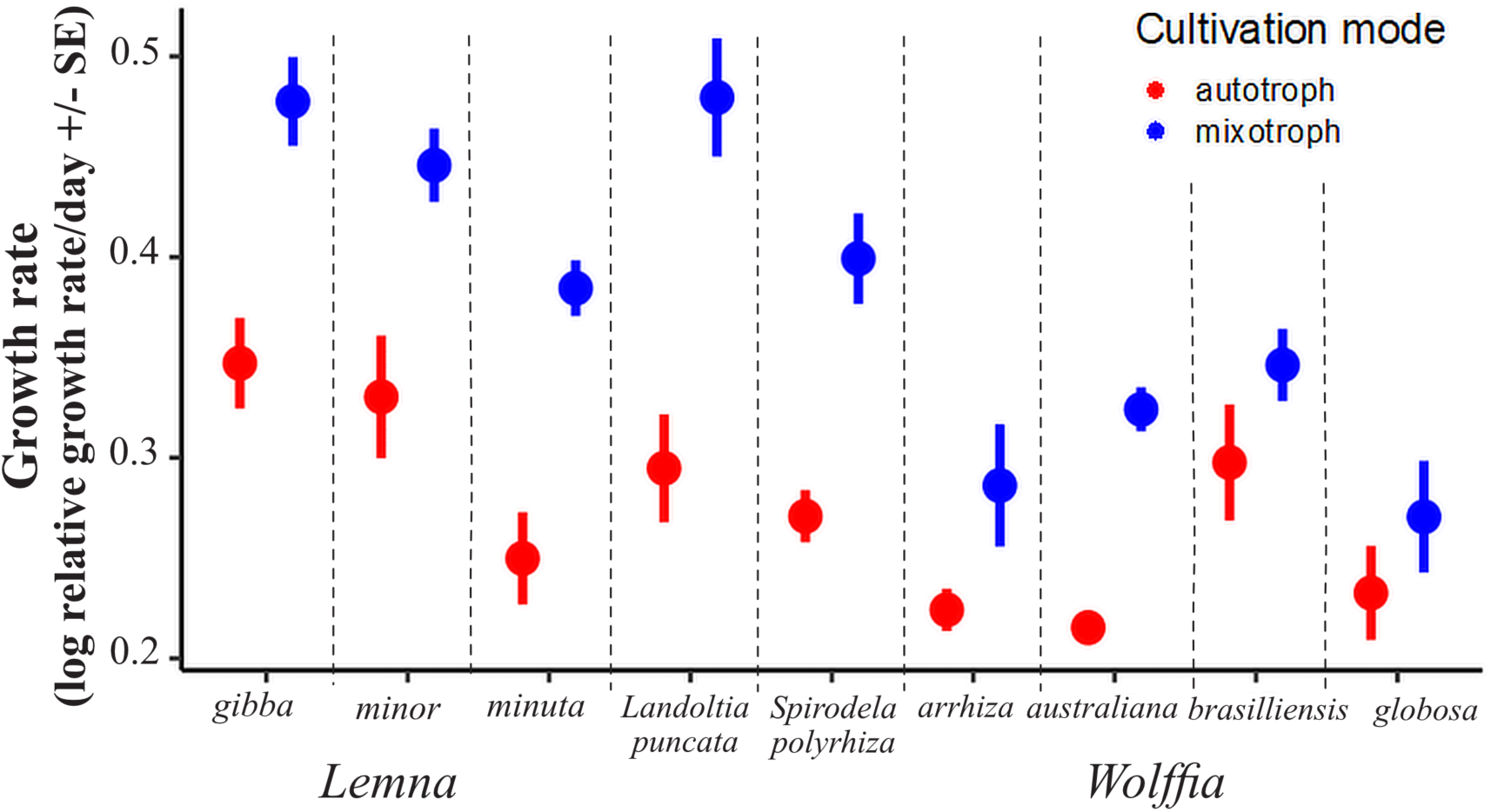
Relative growth rates per day of nine duckweed species, where RGR is the relative increase of the DW per unit time of one day.

### Chlorophyll concentration of duckweed species

Chlorophyll a + b contents of duckweed was normalized to the DW of the plants (Figure 3). Our results suggest that organic carbon supplementation significantly changed chlorophyll content (ANOVA, p < 0.01; Table S2). Furthermore, the chlorophyll content of the rooted duckweed species was higher than that of the rootless species, suggesting higher photosynthetic activities of the rooted species. However, the overall observed differences were not statistically significant (ANOVA, p > 0.01; Table S2).

**Figure 3.**
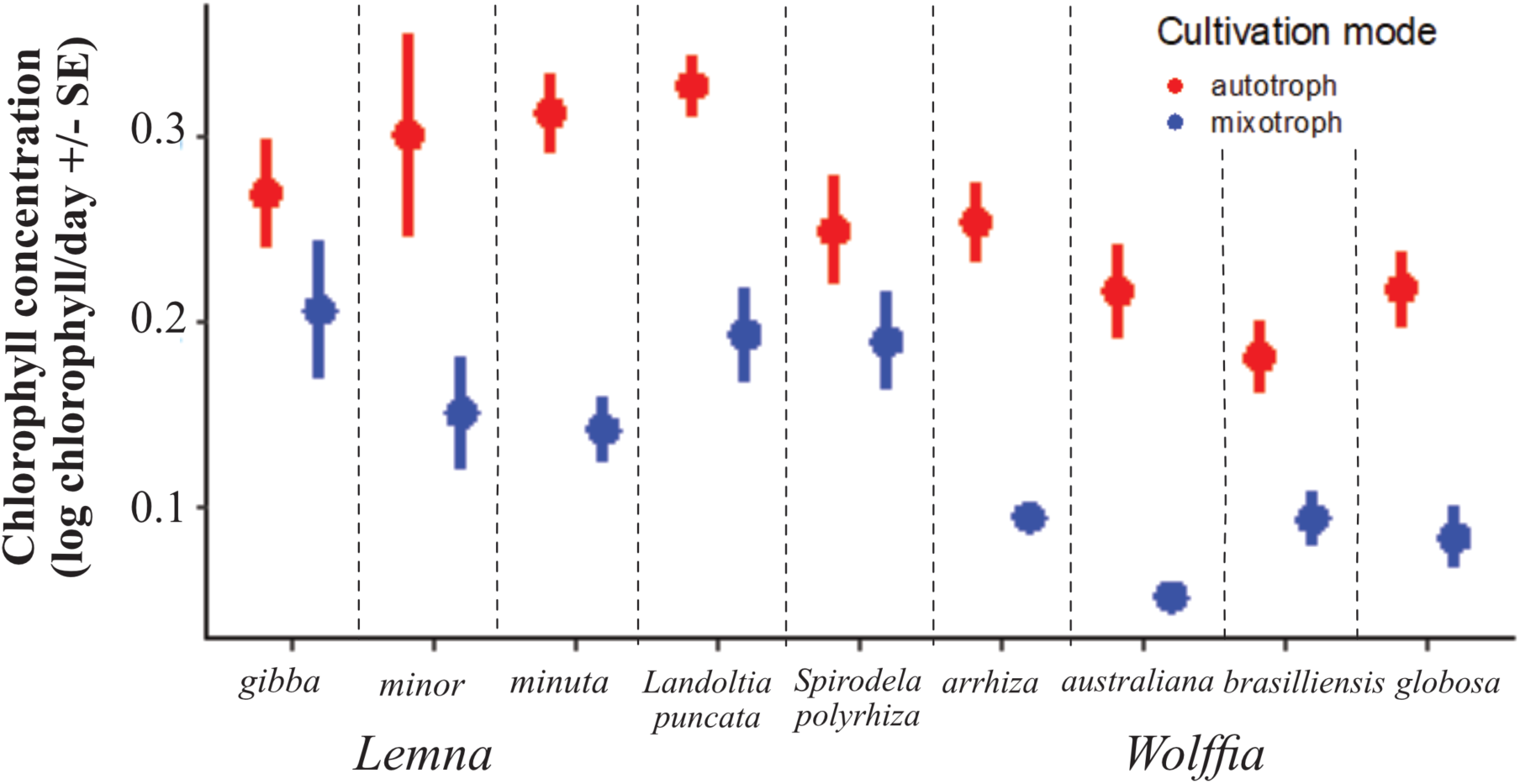
Chlorophyll (a+b) mg^-1^ day^-1^ in various species of duckweed.

### Cobalamin Presence and Concentrations in Duckweeds

To determine whether the presence of bioavailable cobalamin is prevalent among duckweeds or limited to *W. globosa* Mankai ^2^, the nine different species of duckweed were investigated. Furthermore, we explored the effect of trophic conditions on the accumulation of cobalamin in duckweed.

We estimate the presence and content of cobalamin using both biological and analytical methods. Both methods have shown that cobalamin is prevalent in all duckweed species tested (Table 2). To establish whether the detected cobalamin is bioavailable, we differentiated between cyanocobalamin and pseudocobalamin using LC-MS/MS. The commercial pseudococbalamin standard is not available, but this corronoid was identified in Spirulina at high concentration ^33^. Therefore, the extraction of spirulina was used as a positive control to pseudocobalamin(Figure S3). Our results suggest that all species tested contained cyanocobalamin but not pseudocobalamin (Figure S4).

**Table 2:**
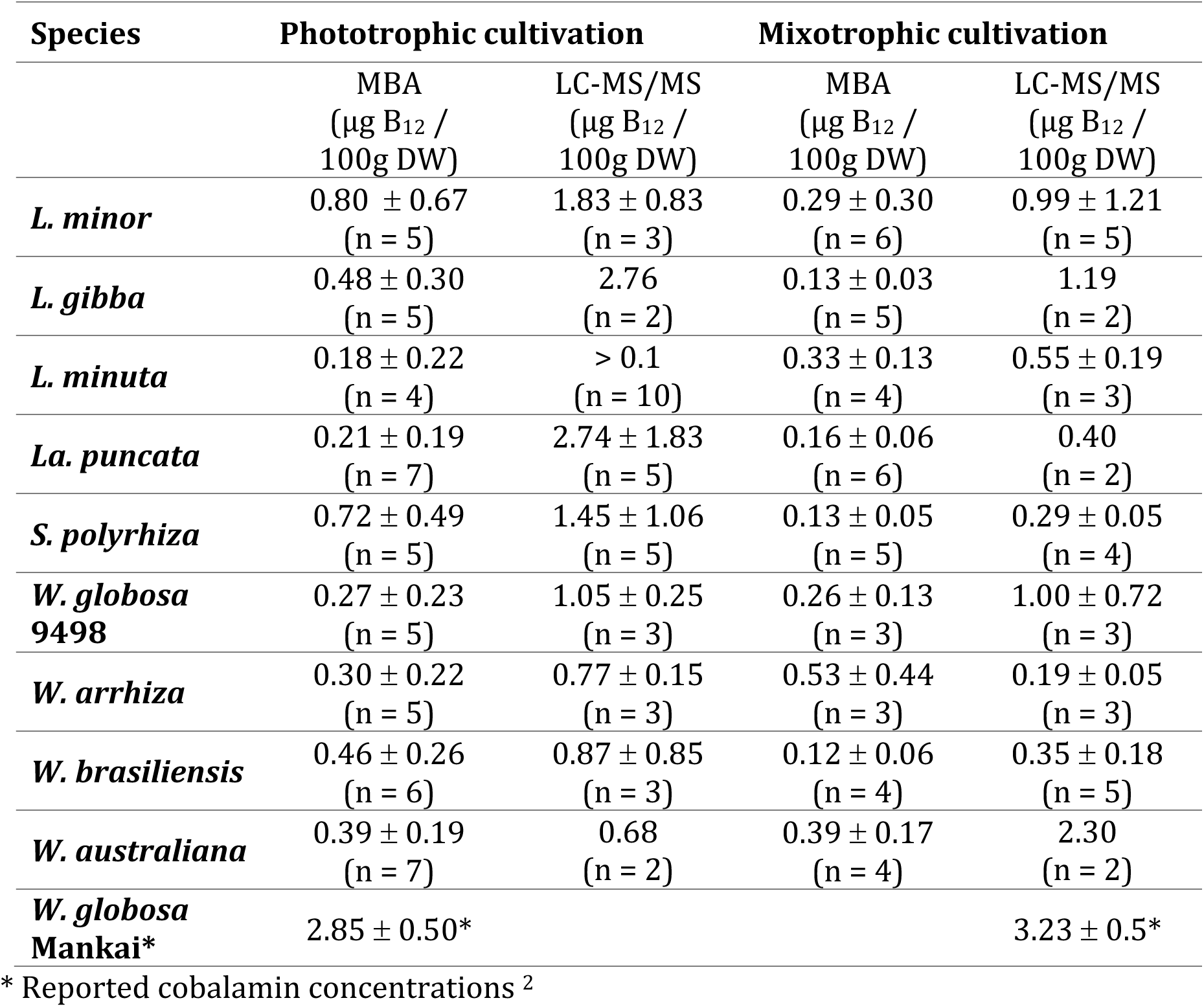
Cobalamin concentrations in duckweed species. Total cobalamin was evaluated using biological (MBA) and analytical (LC-MS/MS) analyzes. The average levels of cobalamin are presented as µg B_12_ per 100 g of duckweed DW.

Peaks are shown at a retention time of 3.8 min and m/z 678.77 confirming the precursor ions for cyanocobalamin, while the presence of peseudocobalamin was not confirmed (Figure S4). Although the m/z was 672.78, match the mass of pseudocobalamin, retention times differed and expected product ions (Figure S3) were not detected in the duckwed extract (Figure S4) suggesting a different molecule.

In particular, the concentrations measured by the bioassay did not necessarily agree with the cobalamin concentrations determined by the analytical method. Generally, the cobalamin values measured by the LC-MS/MS method were two to five times higher than the values estimated by the MBA (Table 2). LC-MS/MS analysis is a direct measurement of the analyte mass, whereas the MBA kit uses the bacterium *L. delbrueckii,* which was certified to estimate cobalamin concertation in food products such as infant formula, cereals, pills, powders, juice, and milk (AOAC-RI). The growth of *L. delbrueckii* could be inhibited by various compounds, such as proteins and fatty acids, that may be present in duckweed extracts (Partanen et al., 2001). Duckweed extracts were shown to be rich in fatty acids and non-polar components, such as carotenoids and phytosterols ^12–14^. These compounds could have interfered with MBA tests with duckweed extracts. Therefore, we suggest that the bioassay should be validated for the plant matrix as the duckweed extract can interfere with the bioassay, yielding cobalamin reads that are lower than certified food products.

The effect of trophic conditions on the accumulation of cobalamin in duckweed species was estimated using MBA and LC-MS/MS (Table 2). The addition of exogenous carbon generally led to lower levels of cobalamin in the duckweeds tested (Table 2, Figure S2). In the presence of organic carbon, cobalamin levels of *L. gibba*, *L. minor*, *La. puncata, S. polyrhiza,* and *W. brasiliensis* were lower than under phototrophic conditions. In contrast, cobalamin levels in *L. minuta* were higher in mixotrophic conditions compared to phototrophic conditions. In *W. globosa* 9498, cobalamin levels were similar under both tropic conditions, while in *W. australiana* and *W. arrhiza,* the cobalamin levels measured by MBA contrasted the measured LC-MS/MS values (Table 2, Table S2). However, the observed differences were not statistically significant (p < 0.01, ANOVA).

## Discussion

We have substantiated that the production of bioavailable cobalamin is not unique to *W. globosa* Mankai but is a shared trait among the nine members of the duckweed family tested here. We further demonstrated that the species contained bioavailable cobalamin and not pseudocobalamin. Furthermore, we have shown that the mixotrophic conditions increased the growth rate of the duckweed species, but reduced the amounts of cobalamin produced compared to the autotrophic conditions.

### Mixotrophic and phototrophic growth and photosynthesis in duckweed

Mixotrophic duckweed cultures grew at least twice as fast as phototrophic cultures (Figure 2), but decreased chlorophyll levels (Figure 3). However, the refreshing of the nutrients did not affect the growth rate or the chlorophyll content (Table S1 and S2). Our results also indicate that the growth medium used in this study (1/2 SH) did not result in nutrient depletion or pH changes that affected growth, in contrast to other media types used for duckweed cultivation ^27, 34^. The growth results were consistent with previous reports showing that the duckweeds *L. minor*, *W. brasiliensis*, *S. punctata,* and *S. polyrhiza* grow faster under light or dark conditions when sucrose was added to medium ^22,35^. However, the addition of sucrose inhibited photosynthesis in *L. perpusila*^36^. Other carbon sources such as galactose, fructose, or sorbitol were not available to *L. minor* and *W. brasiliensis,* but were available to *Spirodela* species ^22,35^. Here, the growth rates and chlorophyl content of *W. globosa* and *W. arrhiza* were slightly higher than those of the other duckweed species used in this study (Figures 2 and 3).

The differences in growth and chlorophyl accumulation among the duckweed species tested here (Figures 2 and 3) were consistent with similar variations reported in duckweeds grown under different trophic conditions ^37–39^. Generally, the RGR of the rooted duckweed species were twice those of rootless species (Figures 2 and 3), in line with previous reports ^26^. Additionally, the rate of development of vegetative propagules was shown to differ between rooted and rootless duckweeds, indicating different reproduction strategies ^40^. Similarly, mixotrophic conditions were shown to increase microalgae biomass by up to 25 times compared to phototrophic conditions ^41^. We further showed that the addition of sucrose substantially increased the RGR of rooted duckweeds and, to a lesser extent, in rootless species (Figures 2; Table S1). The results suggest that the presence of roots in duckweed species enhances exogenous carbon acquisition perhaps by providing additional absorption possibilities directly through the roots and/or through the roots symbionts ^23^. Moreover, *Landoltia* and *Spirodella* fronds have multiple roots, while *Lemna* species have a single root ^42^, yet, in the tested species the number of roots did not affect the RGR (Figure 2; Table S1).

### Mixotrophic and phototrophic cobalamin production in duckweed

In a previous study, bioavailable cobalamin levels were demonstrated in the duckweed species *W. globosa* Mankai ^2,42^. Here, the LC-MS/MS analysis suggested that bioavailable cyanocobalamin was detected in nine duckweed species tested (Table 2, Figure S2), while pseudocobalamin was not identified (Figure S3 and S4). The pseudocobalamin was verified by running a spirulina extract, which was previously shown to contain both cobalamin and pseudocobalamin, ^11,43^. Furthermore, the presence of an exogenous carbon source inhibited the production of cobalamin (Table 2, Figure S2). In contrast, the mixotrophic cultivation of the microalgae *Euglena gracilis* increased the levels of vitamins C and E compared to phototrophic conditions ^44^.

Cobalamin production always depends on a small group of bacteria and archaea that can produce vitamin ^45^. Cobalamin was shown to mediate specific associations within microbial communities ^6^ and play an essential role in molecular interactions that shape microbial communities ^46^. Many algae require cobalamin for growth because their methionine synthase is cobalamin dependent. In this respect they differ from plants, which mainly use cobalamin independent methionine synthase. To obtain cobalamin algae engage in symbiotic relationship with cobalamin producing bacteria and in return supply fixed carbon ^47,48^. Similarly, enteric bacteria are the source of cobalamin in ruminants ^49^. Other eukaryotes, including nematodes and amoeba ^50^, also acquire cobalamin from bacterial producers. The detection of cobalamin in duckweeds raises many questions regarding potential producers and their interactions with host plants and their microbiome. To answer these questions, we currently explore in detail the role of cobalamin in duckweed-microbe interactions to elucidate the pathways and ecological role of cobalamin producers in the duckweed microbiome.

## Conclusions

The recommended transition to healthier and sustainable diets will require a reduction of at least 50% in the consumption of animal-based foods and a 100% increase in plant- based foods ^51^. However, a major disadvantage of plant-based diets is the low levels of bioavailable cobalamin. Only a handful of plant species could provide humans with adequate amounts of bioavailable cobalamin ^10^. Non-animal-derived foods with detectable cobalamin content were mainly restricted to algae and mushrooms, but most of their cobalamin is pseudocobalamin ^10^. However, a recently introduced plant, the *W. globosa* Mankai duckweed, surpasses the bioavailable cobalamin concentrations previously shown in other plants ^2^. Our study showed that the production of bioavailable cobalamin is not unique to Mankai, but prevalent in all nine species of duckweed tested (Table 2), while pseudocobalamin was detected in none of the species (Figure S4). We also showed that cultivation conditions can not only affect duckweed growth rates but also cobalamin production, particularly when duckweed species use an external carbon source (Table 2). We are currently exploring the underlying mechanisms of cobalamin production by duckweed endophytes and their response to changes in trophic conditions.

## Supporting information

Effect of refreshment of nutrients during growth on cobalamin content in the duckweed species (Figure S1); Cobalamin levels in duckweed species cultivated under auto- and mixotrophic conditions (Figure S2); Chromatograms of extraction of *L. minuta* and Spirulina used as reference for pseudocobalamin chromatogram (Figure S3); Chromatograms of *L. minuta* (A), *L. gibba* (B), *L. minor* (C), *La puncata* (D), *S. polyrhiza* (E), *W. australiana* (F), *W. brasiliensis* (G), and *W. globosa* (H) to identify cobalamin types (Figure S4); ANOVA tests for statistical significance of duckweed growth rates under different trophic conditions tested (Table S1); ANOVA tests for statistical significance of duckweed chlorophyll content under different trophic conditions tested (Table S2); ANOVA tests for statistical significance of duckweed vitamin B_12_ content under different trophic conditions tested(Table S3); ANOVA tests for statistical significance of duckweed vitamin B_12_ content under different trophic conditions tested (Table S4); Raw data of cobalamin concentrations in duckweed species (Table S5).

## Acknowledgment

This research was supported by the Israel Ministry of Agriculture and Rural Development Grants No 16-38-0038 and ICA for OG. The Goldinger Trust for OG and IKG also supported this study. The authors acknowledge the ZIWR and AKIS scholarships for their generous support to LK. We gratefully acknowledge the help and support from Prof Appenroth for providing us with the duckweed strains and for his help and support.

## Notes

### Competing Interest Statement

The authors have declared no competing interest.

